# Exploring and explaining properties of motion processing in biological brains using a neural network

**DOI:** 10.1101/2020.09.03.281030

**Authors:** Reuben Rideaux, Andrew E Welchman

## Abstract

Visual motion perception underpins behaviours ranging from navigation to depth perception and grasping. Our limited access to biological systems constrain our understanding of how motion is processed within the brain. Here we explore properties of motion perception in biological systems by training a neural network (‘MotionNet_xy_’) to estimate the velocity image sequences. The network recapitulates key characteristics of motion processing in biological brains, and we use our complete access to its structure explore and understand motion (mis)perception at the computational-, neural-, and perceptual-levels. First, we find that the network recapitulates the biological response to reverse-phi motion in terms of direction. We further find that it overestimates the speed of slow reverse-phi motion while underestimating the speed of fast reverse-phi motion because of the correlation between reverse-phi motion and the spatiotemporal receptive fields tuned to motion in opposite directions. Second, we find that the distribution of spatiotemporal tuning properties in the V1 and MT layers of the network are similar to those observed in biological systems. We then show that compared to MT units tuned to fast speeds, those tuned to slow speeds primarily receive input from V1 units tuned to high spatial frequency and low temporal frequency. Third, we find that there is a positive correlation between the pattern-motion and speed selectivity of MT units. Finally, we show that the network captures human underestimation of low coherence motion stimuli, and that this is due to pooling of noise and signal motion. These findings provide biologically plausible explanations for well-known phenomena, and produce concrete predictions for future psychophysical and neurophysiological experiments.

## INTRODUCTION

The transduction of changing patterns of light into the perception of motion underpins adaptive behaviours ranging from depth estimation to navigation and grasping. In order for motion perception to guide these behaviours effectively, changes in visual input must be translated into accurate estimation of both direction and speed. This – uniquely – requires combining information across space and time. Many biological systems appear to be highly proficient at this task, for example, humans can reliably discriminate differences in speeds between 5-7% (de Bruyn & Orban, 1988; McKee, 1981) and over a century of research on motion processing has expanded our understanding of the neural computations that underlie this ability. However, the biological basis for many aspects of speed estimation remain unknown. A primary constraint on our understanding of these (and other) neural mechanisms is imposed by the limited access we have to biological systems. For example, we can measure the output of the system in response to different inputs (i.e., psychophysics), gross population activity (e.g., fMRI or EEG), or point measurements (i.e., cell recordings), but combining this information to extract the underlying neural computations and principles remains a challenge.

We recently demonstrated the potential of taking an artificial systems approach to bolster understanding of how biological systems function. In particular, we trained a shallow neural network (“MotionNet”) to classify the velocity of motion sequences generated from natural images (Rideaux & Welchman, 2020). Using this approach, we revealed novel relationships between speed and direction encoding, and explained drivers of biases in population tuning and perception. Here we sought to extend this approach to test aspects of motion processing in relation to spatial and temporal frequency characteristics. Moreover, the architecture of the neural network used in our previous study constrained the units in the output layer that we described as being analogous to the middle temporal area (MT) in the primate visual system. This stood in contrast to the layer containing units that is analogous to V1 that was unconstrained and therefore allowed us to gain valuable insights into population characteristics (e.g., tuning biases) that were chosen by the network to best estimate velocity. In this paper we used a new neural network that did not predefine the V1 of MT stages of the model. Specifically, we train a new neural network (“MotionNet_xy_”) to estimate continuous measures of horizontal and vertical velocity by including an additional regression layer. This does not constrain the properties of the MT layer units, allowing them to develop characteristics that best serve the task of velocity estimation.

Using this artificial systems approach we examine how spatiotemporal information is combined to produce our (mis-)perceptions of image velocity. For instance, when image contrast is reversed the between motion frames this produces a corresponding reversal in perceived motion direction (Anstis, 1970). Electrophysiological work shows that this perceptual illusion is also reflected in the responses of macaque V1 and MT neurons: their preferred direction is inverted (Duijnhouwer & Krekelberg, 2016). After verifying this behaviour in the artificial system, we explore how these changes influence the calculation of speed. Unpublished observations suggest that observers over and underestimate the speed of slow and fast reverse-phi motion, respectively (Parthasarathy, 2019; Ruda, Riesen, & Hock, 2016). We find that the network exhibits the same biases, and then use our access to the system to show that this due to the similarity between reverse-phi motion and the receptive fields of spatiotemporal neurons tuned to opposite directions.

We then examine how spatial and temporal information is combined to compute speed. Electrophysiological work shows that V1 neurons are tuned to a range of spatial and temporal frequencies (Friend & Baker, 1993; Holub & Morton-Gibson, 1981; Tolhurst & Movshon, 1975), but their tuning for these properties are independent. By contrast, MT some neurons appear to show speed tuning, requiring joint encoding of spatial and temporal frequency (Perrone & Thiele, 2001; Priebe, Cassanello, & Lisberger, 2003). It has been proposed that MT neurons tuned to slow speeds receive input from V1 neurons sensitive to high and low spatial and temporal frequencies, respectively, while the opposite is true for MT neurons tuned to high speeds. This notion is supported by some neurophysiological evidence (Priebe et al., 2003), but remains a challenge to directly test in biological systems due to the difficulty of tracking synaptic connections between brain regions. By contrast, the connections between layers in the artificial system are equally accessible as all its architecture; thus, we test this possibility and find that the relationship predicted between spatiotemporal V1 and MT neurons in biological systems is evident in the network.

Although some MT neurons appear achieve speed selectivity by pooling V1 activity, neurophysiological work suggests that many MT neurons exhibit selectivity indistinguishable from V1 neurons, i.e., separable tuning to spatial and temporal frequency (Priebe et al., 2003). Similar diversity across MT neurons is also observed for direction selectivity, i.e., whether a neuron responds to the individual components or combined pattern of a moving object (Movshon, Adelson, Gizzi, & Newsome, 1986). These two properties index the complexity of the information that is encoded by MT neurons in terms of speed and direction, and we find that they are positively correlated among in the network, that is, MT units tuned to speed are more likely to be also tuned to pattern motion. Despite this, we find that MT neurons tuned to less complex features are more influential in the final velocity estimation.

Finally, we show that the network recapitulates neural and psychophysical performance in response to reduced motion coherence (Britten, Shadlen, Newsome, & Movshon, 1992), exhibiting the same speed opponency, noise reduction, mechanisms observed in biological systems (Mikami, Newsome, & Wurtz, 1986). In particular, we show that MotionNet_xy_ underestimates the speed of low coherence motion stimuli (Schütz, Braun, Movshon, & Gegenfurtner, 2010) and demonstrate that this is due to pooing of noise and signal motion.

## METHOD

### Naturalistic motion sequences

To train a neural network to estimate image velocity, we generated motion sequences using 200 photographs from the Berkeley Segmentation Dataset (https://www2.eecs.berkeley.edu/Research/Projects/CS/vision/bsds/). Images were greyscale indoor and outdoor scenes (converted from RGB using MATLAB’s (The MathWorks, Inc., Matick, MA) *rgb2grey* function). Motion sequences (six frames) were produced by translating a 32 × 32 pixel cropped patch of the image (**Fig. 1a**). Motion direction and speed were randomly assigned from uniform distributions between 0-360° and 0.8-3.8 pixels/frame, respectively. Images were translated in polar coordinates, e.g., an image moving at a speed of 1 pixel/frame in 0° (right) direction was translated by +[x=1,y=0] per frame, while an image moving at the same speed in 45° direction was translated +[x=.7071,y=.7071]. Image translation was performed in MATAB using Psychtoolbox v3.0.11 subpixel rendering extensions (Brainard, 1997; Pelli, 1997) (http://psychtoolbox.org/). The speeds used to train the network were selected because they did not exceed the image dimensions (32 × 32 pixels) and matched those used in our previous study (Rideaux & Welchman, 2020). We generated 32,000 motion sequences, which were scaled so that pixel intensities were between –1 and 1, and randomly divided into training and test sets, as described in the Training Procedure section.

**Figure 1.**
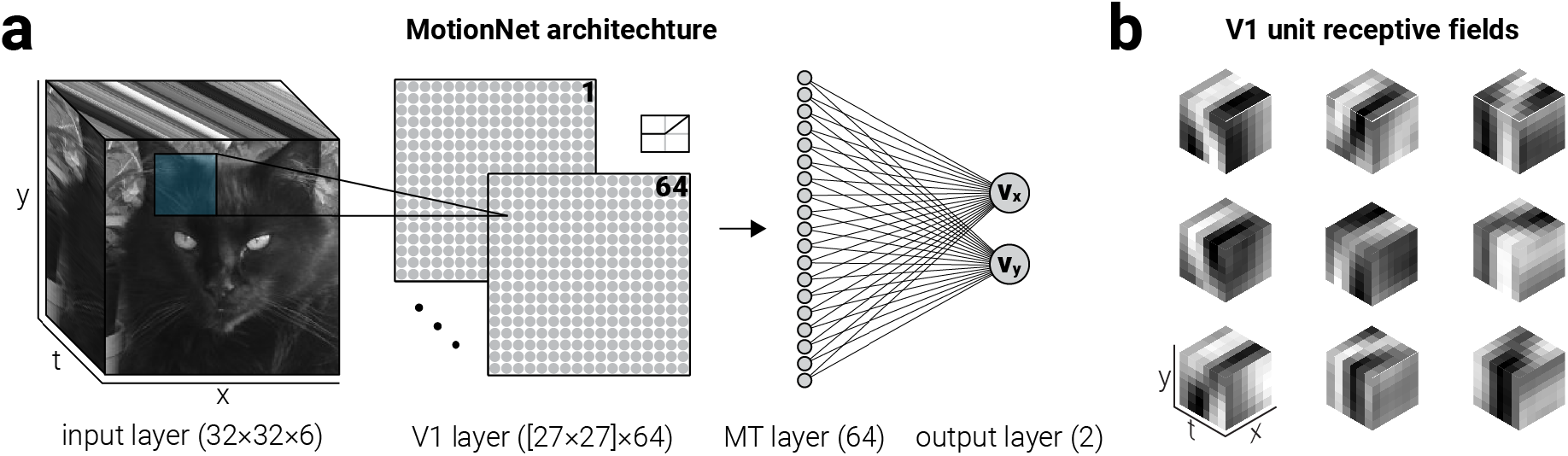
MotionNet_xy_ architecture. (**a**) MotionNet_xy_ was initialized with an input layer, convolutional and dense layers representing V1 and MT, respectively, and an output regression layer. (**b**) Following training on motion sequences, kernels (V1 units) that were initialized as Gaussian noise formed three-dimensional Gabors; nine examples, selected at random, are shown

### MotionNet_xy_ architechture

All the networks described in the study were implemented in Python v.3.6.4 (https://python.org) using Tensorflow (www.tensorflow.org), a library for efficient optimization of mathematical expressions. We used a convolutional neural network that comprised (i) an input layer, one convolutional-pooling layer, (iii) one dense layer, and (iv) an output regression layer (**Fig. 1a**).

Inputs were image patches (32 × 32 × 6 pixels; the last dimension indexing the motion frames). In the convolutional layer, inputs passed through 64 three-dimensional kernels (6 × 6 × 6 pixels) producing 64 two-dimensional output maps (27 × 27 pixels). This resulted in 18,496 units (64 maps of 27 × 27 pixels) forming 10,077,696 connections to the input layer (64, 27 × 27 × 6 × 6 × 6 pixels). Since mapping is convolutional, this required that 13,888 parameters were learned for this layer (64 filters of dimensions 6 × 6 × 6 plus 64 offset terms). We chose units with rectified linear activation functions to model neurophysiological data (Movshon, Thompson, & Tolhurst, 1978). The activity, *a*, of unit *j* in the *k*^th^ convolutional map was given by:

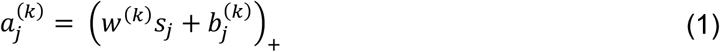

where *w*^(*k*)^ is the 6 × 6 × 6 dimensional 3D kernel of the *k*^th^ convolutional map, *s*_j_ is the 6 × 6 × 6 motion sequence captured by the *j*^th^ unit, *b*_j_ is an offset term and (.)_+_ denotes a linear rectification non-linearity (ReLU). Parameterizing the motion image frames separately, the activity 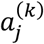 can be alternatively written as:

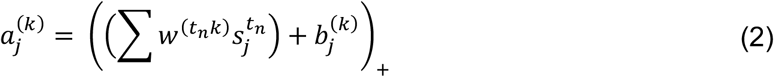

where *w*^(*t_n_k*)^ represent the *k*^th^ kernels applied to motion image frames (i.e., receptive fields at times 1 to 6), while 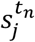 represent the input images captured by the receptive field of unit *j*.

A dense layer (1,183,776 connections; 23,328 per feature map, resulting in 1,183,744 parameters including the 64 offset terms) mapped the activities in the pooling layer to 64 fully connected units. The vector of dense layer activities *r* was obtained by mapping the vector of activities in the convolutional layer a via the weight matrix *W* and adding the offset terms *b*:

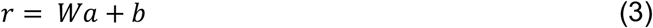

Finally, a regression layer (128 connections, 64 for each of the two regression units, resulting in 130 parameters including the 2 offset terms) mapped activities from the dense layer to two regression units, which represented the x and y velocity of the motion sequence. The regression unit activities were obtained using equation (3).

### Training procedure

Motion sequences were randomly divided into training- (75%, *n*=24,000) and test- (25%, *n*=8,000) sets. No sequences were simultaneously present in the training and test sets. To optimize MotionNet_xy_, only the training set was used. We initialized the weights of the convolutional layer as Gaussian noise (mean, 0; s.d., 0.001). The weights in the dense and regression layers and all offset terms were initialized to zero.

MotionNet_xy_ was trained using mini-batch gradient descent with each batch comprising 32 randomly selected examples. For each batch, we computed the derivative of the mean squared loss function with respect to parameters of the network via back-propagation, and adjusted the parameters for the next iteration accorded to the update rule:

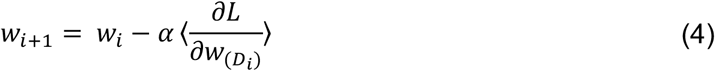

where *α* is the learning rate, and 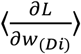 is the average over the batch *D_i_* of the derivative of the loss function with respect to the *w*, evaluated at *w_i_*. The learning rate *α* was constant and equal to 1.0e^-4^. After evaluating all the batches once (i.e., completing one epoch) we tested MotionNet_xy_ using the test image dataset. We repeated this for 25 epochs.

### Generation of test stimuli

A range of stimuli were used to test the response of the network after it had been trained on natural images. With the exception of sinewave and plaid stimuli, which were generated in Python using in-house scripts, all stimuli were generated using the Python toolbox Psychopy (Peirce, 2007) v1.90.3 (http://www.psychopy.org).

### Decoding direction and speed

To avoid issues associated with using a circular variable to train a regression output, the network was trained to estimate the x and y velocity of motion sequences. These estimates were then converted to speed *ρ* and direction *ϕ* with the following:

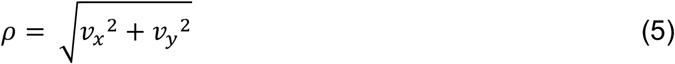

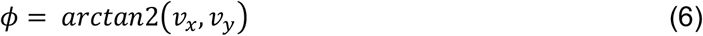

where *v_x_* and *v_y_* denote x and y velocity vectors.

### Component- and pattern- motion selectivity

To compare the component- and pattern-motion selectivity of MotionNet_xy_ units to those of neurons in macaque V1 and MT (extracted and replotted neurophysiological data from Figures 11-13 of (Movshon et al., 1986)), we measured the activity of V1/MT units in response to sinewave gratings and plaids (135° separation) moving in 16 evenly spaced directions between 0 and 360° at its preferred spatial- and temporal-frequency (**Fig. 2c**).

**Figure 2.**
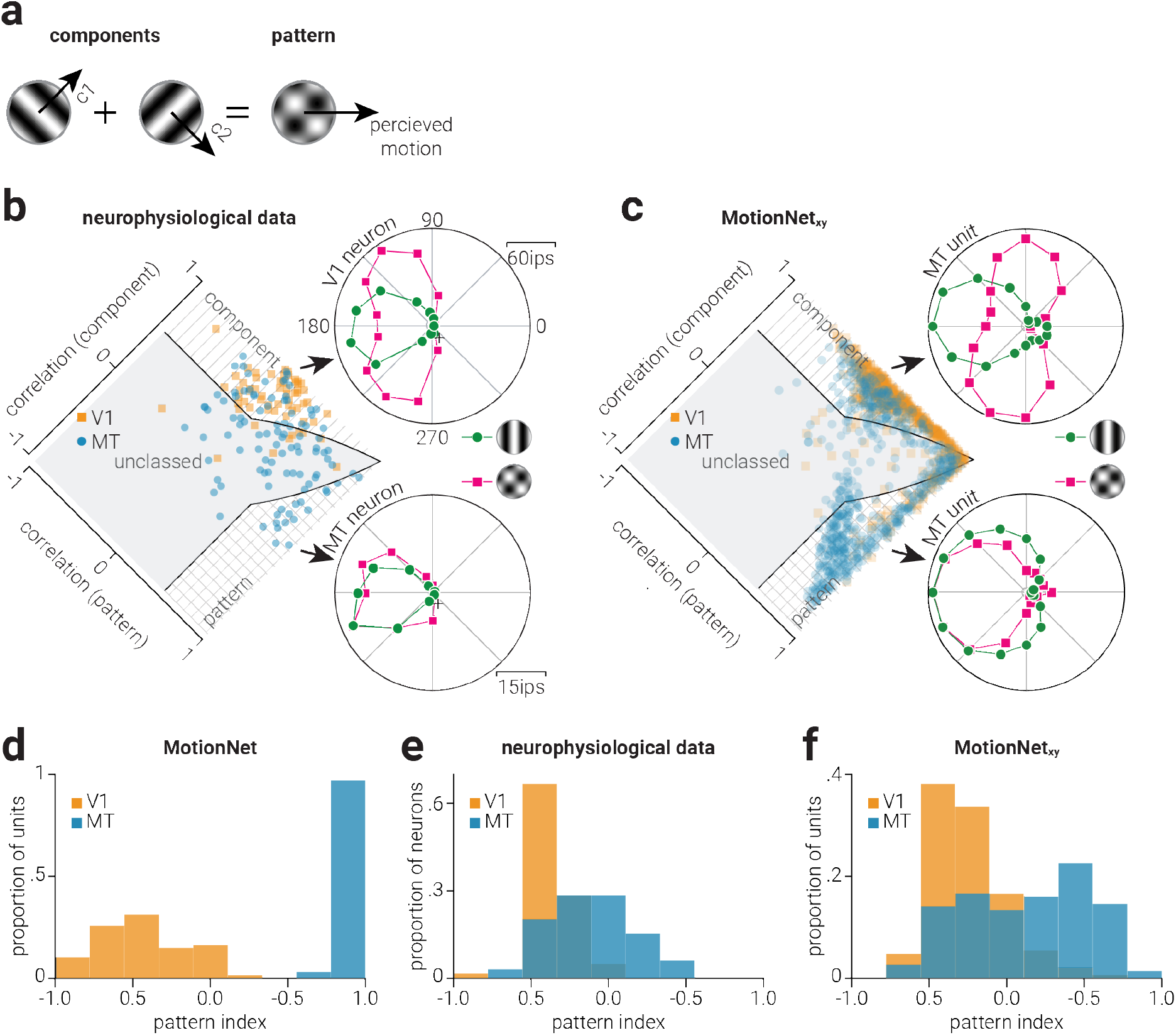
Biological and artificial visual system direction selectivity. (**a**) Illustration of how two “component” sinewave gratings moving in different directions form a plaid “pattern” which moves in a (different) third direction. (**b**) Data from (Movshon et al., 1986) showing single neuron responses in V1 (top) and MT (bottom) to a sinewave grating vs. a plaid stimulus. The distribution plot shows the population of single neuron responses, and whether they are classified as component-motion or pattern-motion selective. (**c**) The same as (**b**), but for MotionNet_xy_; single polar plots (top and bottom) both show responses of MT units classified as either component- or pattern-motion selective. (**d**) Proportion of the previous “MotionNet” (Rideaux & Welchman, 2020) V1 and MT units as a function of pattern index. (**e-f**) Same as (**d**), but for neurophysiological data (Movshon et al., 1986) and the new MotionNet_xy_ network.

To classify each unit as component-selective (i.e., selective for the motion of the individual components comprising a plaid pattern), pattern-selective (i.e., selective for the motion of the plaid pattern), or unclassed (**Fig. 2c**), we used the method described in (Movshon et al., 1986). Briefly, we compared the unit responses to ideal ‘component’ and ‘pattern’ selectivity using goodness of fit statistics. As the component and pattern selectivity responses may be correlated, we used the partial correlation in the form:

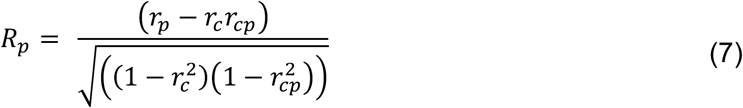

where *R_p_* denotes the partial correlation for the pattern prediction, *r_p_* is the correlation of the data with the pattern prediction, *r_c_* is the correlation of the data with the component prediction, and *r_cp_* is the correlation of the between the two predictions. The partial correlation for the component prediction was calculated by exchanging *r_c_* for *r_p_* and vice versa. We labelled units as “component” if the component correlation coefficient significantly exceeded either zero or the pattern correlation coefficient, whichever was larger. Similarly, we labelled units as “pattern” if the pattern correlation coefficient significantly exceeded either zero or the component correlation coefficient. Units were labelled as “unclassed” if either (i) both pattern and component correlations significantly exceed zero, but do not differ significantly from one another, or (ii) neither correlation coefficient differed significantly from zero. To demonstrate the consistency in training outcomes, we trained ten networks and in **Figure 2** present the cumulative distribution of all 10 networks.

To compare the distribution of pattern-motion selectivity among V1 and MT units in MotionNet_xy_ with those of our previous network (“MotionNet”; Rideaux & Welchman, 2020) and V1 and MT neurons, we projected the values shown in **Figure 2b** and **Figure 2c**, in addition to data from Figure 3e our previous study (Rideaux & Welchman, 2020) along the diagonal to establish a unified estimate of pattern-motion selectivity for each unit (**Fig. 2d-f**).

### Reverse-phi motion responses

To compare the phi and reverse-phi responses of MotionNet_xy_ units to those of neurons in macaque V1 and MT (extracted and replotted neurophysiological data from Figures 3a and 4a of (Duijnhouwer & Krekelberg, 2016)), we measured the activity of V1/MT units in response to dot motion. Dot motion stimuli in the phi condition consisted of 5 randomly positioned white dots (pixel value, 1.0; radius, 4 pixels) on a mid-grey background (pixel value, 0.0), which were allowed to overlap (with occlusion) and wrapped around the image when their position exceeded the edge. Of the six motion sequence frames presented, only the first two frames comprised dot motion, while the last four were presented as uniform mid-grey. For each V1/MT unit, we presented dot motion stimuli moving in 16 evenly spaced directions (0-360°), at their preferred speed. The reverse-phi dot motion stimuli were the same as those used in the phi condition, except the contrast of the dots was reversed (from white to black) on the second frame. The responses of V1 and MT units from ten networks were aligned to a common preferred direction and the average for each are shown in **Figure 3b-c**.

To test how MotionNet_xy_ estimated the speed of reverse-phi stimuli, we compared the speed decoded by the network in response to the phi and reverse-phi stimuli described above over a range of speeds (five linearly spaced speeds between 1.0 and 3.5 pixels/frame). We tested ten networks and the average and standard deviation of their estimated speed is shown in **Figure 3d**. To explore why MotionNet_xy_ misjudges the speed of reverse-phi stimuli, we separated the V1 and MT units in two groups, those that were more tuned to the displacement direction and those that were more tuned to the opposite-to-displacement direction, by assessing whether they were positively or negatively weighted to the v_x_ regression output unit, respectively. This classification was straightforward for MT units, which are directly connected to the regression layer, but for V1 units we used the classification of the MT unit for which each V1 unit was most positively weighted. We then measured the average activity of these subpopulations of V1 and MT units in response to the phi and reverse-phi stimuli. Finally, to explain why the speed of reverse-phi motion is misjudged, we ran a simulation on a simplified version of the phenomena. The simulation consisted of computing the cross-correlation between phi and reverse-phi stimuli (16 × 16 × 2 [x,y,t] pixel image sequence comprising a white [pixel value, 1] and black [pixel value, −1] vertical edge centred on the midline at time 0, and moving at one of 3 displacements speeds (1, 2, and 3 pixels) to the right (+v_x_) at time 1) and a bank of four spatiotemporal filters (8 × 8 × 2 [x,y,t] pixels comprising a white and black vertical edge centred on the midline at time 0 and moving at the same displacement speed as the phi/reverse-phi stimuli to the right (+v_x_) or to the left (-v_x_) at time 1). The reverse phi stimulus was the same as the phi stimulus, except that it reversed polarity at time 1, and both combinations of light-dark and dark-light edge filters were used. For each cross-correlation we calculated the average of value. To emulate the computations of MotionNet_xy_, only positive and valid cross-correlation values were included.

### Spatiotemporal tuning properties

To compare the properties of V1 and MT units that emerged within MotionNet_xy_ to those of V1 and MT neurons in biological systems, we extracted neurophysiological data of owl monkey V1 neurons from Figure 9A and Figure 10A of (O’Keefe, Levitt, Kiper, Shapley, & Movshon, 1998) and re-analysed data of macaque MT neurons from (Wang & Movshon, 2016). To establish the spatial and temporal frequency tuning preferences of MotionNet_xy_ V1 and MT units we tested the network with drifting sinewave gratings. The direction and spatiotemporal tuning preference of each unit was determined as the stimulus movement direction, spatial frequency, and temporal frequency that produced maximal activity (**Fig. 4a-c**, right). Sixteen directions (linearly spaced between 0-360°), ten spatial frequencies (logarithmically spaced between 8 and 25 pixels/cycle), and ten temporal frequencies (logarithmically spaced between 4 and 25 cycles/frame) were tested, resulting in 1600 (16 × 10 × 10) stimulus types. For each stimulus type, we computed the average activation of 32 gratings at evenly spaced starting phase positions between 0 and 360°.

To assess the input from the V1 layer to MT units tuned to different speeds, we first established the preferred speed of MT units *ρ_MT_* with:

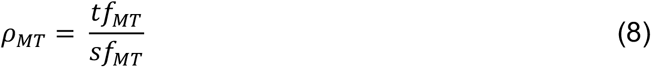

where *sf_MT_* and *tf_MT_* denote the preferred spatial and temporal frequency of the MT unit. Then, for each V1 unit, we established the MT unit to which it was maximally connected and used a median split to separate the V1 units into those maximally connected to MT units that preferred slower or faster speeds. Finally, we compared the preferred spatial and temporal frequency tuning of these distributions (**Fig. 4d-e**). To demonstrate the consistency in training outcomes, we trained ten networks and in **Figure 4** present the mean values with error bars showing s.d.

### Separable and covariate spatiotemporal tuning properties

To compare the separable spatial/temporal-frequency and speed-selectivity of MotionNet_xy_’s units to those of neurons in macaque MT (extracted and replotted neurophysiological data from Figures 5b-d of (Priebe et al., 2003)), we measured the activity of V1/MT units in response to sinewave gratings moving in their preferred direction at from 12 spatial frequencies (logarithmically spaced between 8 and 25 pixels/cycle), and 12 temporal frequencies (logarithmically spaced between 4 and 25 cycles/frame), resulting in 144 (12 × 12) stimulus types. This method yielded spectral responses maps for each V1/MT unit in the network. We used the method described by Perrone & Thiele (2001) to fit a twodimensional Gaussian model to the spectral response maps according to the following equation:

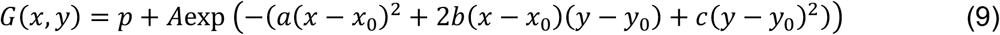

where *G*(*x,y*) denotes the unit response at location (*x,y*), *p* is a constant offset, *A* is the amplitude of the peak, (*x*_0_, *y*_0_) is the location of the center of the peak, and *a, b*, and *c* are positive-definite and defined as

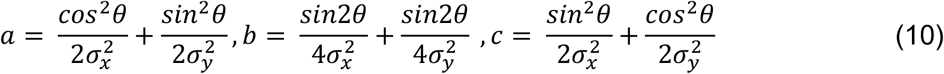

where *θ* denotes the orientation of the peak, and *σ_x_* and *σ_y_* indicate the width of the peak in x and y dimensions, respectively. To classify the units as independently tuned to spatial- /temporal frequency, speed tuned, or unclassified, we used the method described by (Priebe et al., 2003); that is, we compared the correlation of the each unit’s spectral response map to the model fit described in equation (9) where the orientation is either zero (independent tuning) or at an angle that aligns the peak to the origin (speed tuning). Using these values, we performed the same assay as was conducted to determine the component- and pattern-motion selectivity to establish their independent and speed selectivity (**Fig. 5d**).

To compare the distribution of speed selectivity among MT units in MotionNet_xy_ to that among MT neurons we projected the values shown in **Figure 5a** and **Figure 5b** along the diagonal to establish a unified estimate of speed selectivity for each unit (**Fig. 5c,d**). To assess the relationship between pattern-motion and speed selectivity of MotionNet_xy_ units we computed the Pearson correlation between pattern and speed indices of MT units (**Fig. 5e**). In line with previous neurophysiological work (Priebe et al., 2003), units that were unclassified in both dimensions were omitted from the correlation analysis. To demonstrate the consistency in training outcomes, we trained ten networks and in **Figure 5** present the values of all ten networks.

### Speed opponency

To compare the direction discrimination performance of MotionNet_xy_ at varying levels of motion coherence to neurophysiological recordings from macaque (extracted and replotted neurophysiological data from Figures 9a and 11a of (Mikami et al., 1986)), we measured individual MT unit activity in response to dot motion stimuli (dot pixel value, 1.0; background pixel value, −1.0; dot radius, 4 pixels) moving in either the preferred or nonpreferred direction at eight logarithmically (base 2) spaced speeds between the minimum (0.8 pixels/frame) and maximum (3.8 pixels/frame) speeds used to train the network.

### Motion coherence

To compare the direction discrimination performance of MotionNet_xy_ at varying levels of motion coherence to neurophysiological and psychophysical recordings from macaque (extracted and replotted neurophysiological/psychophysical data from Figures 4 and 6 of (Britten et al., 1992)), we measured the direction estimates of the network in response to dot motion stimuli. Dot motion stimuli consisted of 333 randomly positioned white dots (pixel value, 1.0; radius, 2 pixels) on a black background (pixel value, −1.0), which were allowed to overlap (with occlusion) and wrapped around the image when their position exceeded the edge. A proportion of the dots moved in the signal direction, while the remaining dots moved in directions randomly sampled from 0 to 360°; all dots moved at 3 pixels/frame. Seven coherence levels were tested, logarithmically spaced between 0.001 to 0.2. For each coherence level, 100 trials were performed and estimates within ±90° of the signal direction were considered correct. In line with (Britten et al., 1992), we fit a Weibull function to the mean performance to estimate the threshold. Using a similar approach, we compared the speed estimates of MotionNet_xy_ at varying levels of motion coherence with psychophysical data from humans (extracted and replotted psychophysical data from Figures 8b of (Schütz et al., 2010)). For this test, dot motion stimuli consisted of ten randomly position dots, and we used five linearly-spaced coherence levels between 0.2 and 1.0. To test if the MotionNet_xy_ underestimated the speed of partially coherent dot motion stimuli because of pooling noise and signal, we computed the Pearson correlation between the mean activity of MT units across 10 networks in response to 0% and 100% noise, with the activity in response to 50% noise.

### Data reanalysis

Data in **Figures 2b, 2e, 3b-c, 4a-d, 5a, 5c, 7b-d** were extracted from published papers (Britten et al., 1992; Duijnhouwer & Krekelberg, 2016; Mikami et al., 1986; Movshon et al., 1986; O’Keefe et al., 1998; Priebe et al., 2003; Schütz et al., 2010) using WebPlotDigitalizer. Data in **Figure 4c-d** are a reanalysis of archived data (https://archive.nyu.edu/handle/2451/34281) from the published paper (Wang & Movshon, 2016).

### Data availability

We performed analyses in Python using standard packages for numeric and scientific computing. All the code and data used for model optimization, implementations of the optimization procedure, and behavioural data is freely and openly available at repository.cam.ac.uk/XXX (awaiting doi).

## RESULTS

### Network architecture and training

We created an artificial system, which we refer to as “MotionNet_xy_”, tasked with decoding the velocity of image sequences (**Fig. 1a**). The network input comprised a sequence of image frames (*x-y*) depicting a scene moving through time (*t*). This was convolved with three-dimensional kernels (*x-y-t*). The resultant activity was then passed to a dense layer of units. Finally, the activity of the dense layer was read out by two output units, to produce estimates of horizontal (v_x_) and vertical velocity (v_y_). We referred to the convolutional and subsequent dense layer as V1 and MT, respectively, as their hierarchy was analogous to their namesake in biological systems.

We trained MotionNet_xy_ to decode the velocity of natural images moving at a range of speeds (0.8–3.8 pixels/frame) and directions (0-360°). Following training, there was a high correlation between the network’s estimates and the velocity of novel motion sequences (v_x_, *r*=.89; v_y_, *r*=.93). V1 units were initialized with Gaussian noise, but after training they resembled (**Fig. 1b**) receptive fields in primary visual cortex (Movshon et al., 1978; Rust, Schwartz, Movshon, & Simoncelli, 2005).

### Component- and pattern-motion selectivity

To judge an object’s movement, motion signals must be integrated across the stimulus as local motions are often ambiguous (“the aperture problem”). Experimental tests of motion integration often use plaid patterns composed of two sinewave components (**Fig. 2a**). The individual components can move in different directions from the overall plaid (Movshon et al., 1986) and V1 neurons signal motion of the components (Gizzi, Katz, Schumer, & Movshon, 1990; Movshon et al., 1986). For example, the V1 neuron shown in **Figure 2b** responds most strongly to a leftwards moving grating; but when shown a plaid, it responds most strongly to motion above or below leftwards such that one of the component gratings moves leftwards. By contrast, some MT neurons show pattern-motion selectivity (**Fig. 2b**, bottom) – responding to the plaid’s features, rather than the individual components. The response of a neuron to sinewave and plaid stimuli can be used to classify it as either component- or pattern-motion selective. Applying this classification to a population of neurons shows that V1 neurons are exclusively component-motion selective, whereas MT contains a mixture of neurons selective to component- and pattern-motion (**Fig. 2b**). We applied the same analysis to the units of MotionNet_xy_ and found a similar pattern of results (**Fig. 2c**).

We previously showed a similar pattern of selectivity emerged in a neural network (“MotionNet”) trained to make discrete velocity classifications (Rideaux & Welchman, 2020); however, these results differed from biological findings in that MT units were exclusively pattern-motion selective (rather than containing a mixture of selectivity; **Fig. 2d**). This is likely because in the previous network, which performed discrete velocity classifications, MT units were constrained to represent specific velocities. By comparison, units in the MT layer of this network, like the units in V1, were unconstrained and could form characteristics that best served the output regression layer. As a result, here we found a pattern of selectivity that more closely resembled that found in biological systems (**Fig. 2e**): V1 units were component-motion selective while units in the MT layer had a mixture of component- and pattern-selectivity (**Fig. 2f**).

The direction selectivity of neurons can be dramatically altered, as in the case of ‘reverse-phi’ motion, in which the contrast of images in a sequence is reversed between frames (**Fig. 3a**). Perceptually this leads to the impression of movement in the opposite direction from true movement (Anstis, 1970). It has been shown that neurons in V1 and MT will exhibit inverted preferences in this situation, such they respond maximally to reverse-phi stimuli moving in the non-preferred direction (Duijnhouwer & Krekelberg, 2016; **Fig. 3b-c**, left). We found that the activity of MotionNet_xy_’s V1 and MT units were similarly reversed in response to reverse-phi stimuli (**Fig. 3b-c**, right). It is encouraging to see the network recapitulates this well-known phenomenon, but how does it estimate the speed of these stimuli? We tested the network with phi and reverse-phi motion stimuli over a range of displacement speeds. We found that for phi motion, the network consistently underestimated the speed of stimuli, which is likely because the network was trained on motion sequences comprising six frames, whereas our phi stimuli comprised only two (**Fig. 3d**, cyan markers). By contrast, we found that the speed of reverse-phi stimuli was overestimated for low displacement speeds and underestimated for high speeds (**Fig. 3d**, orange markers). Some evidence for the same pattern of behaviour in humans has previously been found (Parthasarathy, 2019; Ruda et al., 2016), but more work is needed to explicitly investigate this phenomenon in biological systems.

To understand why this phenomenon occurs in the network, we measured the activity of V1 and MT units tuned to either the displacement (+v_x_) or opposite-to-displacement direction (-v_x_), in response to phi and reverse-phi motion at different speeds (**Fig. 3e-f**). For phi motion, the activity of the V1 +v_x_ subpopulation stays approximately the same as speed as increased, while that of V1 -v_x_ subpopulation is reduced. This increasing difference in activity between subpopulations of V1 units is propagated to the MT units to produce a divergent pattern of activity. As the difference between subpopulations responses increases, the balance of activity shifts towards the displacement direction, evoking a faster estimate of speed in this direction (**Fig. 3d**, cyan markers). This pattern of responses is consistent with our previous work (Rideaux & Welchman, 2020), where we showed that low speed motion sequences moving in different directions are highly correlated, thus directions are less distinguishable than high speed sequences.

The responses evoked by reverse-phi are markedly different. First, as expected from evidence of the reversal of direction selectivity, the V1 -v_x_ subpopulation are more active than the V1 +v_x_ subpopulation. Second, the activity of both V1 subpopulations is lower than seen for phi motion at the slowest speed and increases with displacement speed. This reflects the evolutionary adaptation of receptive fields to frequently occurring (phi) motion compared with infrequent (reverse-phi) motion. Finally, both subpopulations increase at approximately the same rate, so the relative difference between their activity reduces with displacement speed. To explain why this occurs, we simulated a simplified version of the phenomenon in which we measure the cross-correlation between a phi and a reverse-phi edge stimulus at three displacement speeds with four spatiotemporal filters tuned to leftward and rightward displacement with either light-dark or dark-light polarity arrangement (**Fig. 3g**, left). At the lowest displacement speed (v_x_=1), the cross-correlation for reverse-phi is both attenuated and reversed compared to the cross-correlation for phi (**Fig. 3g**, right). However, the relative difference between the cross-correlation for -v_x_ and +v_x_ filters is larger for reverse-phi. With increasing displacement (v_x_=2 and v_x_=3), the relative difference between -v_x_ and +v_x_ filters increases for phi, while decreasing for reverse-phi.

**Figure 3.**
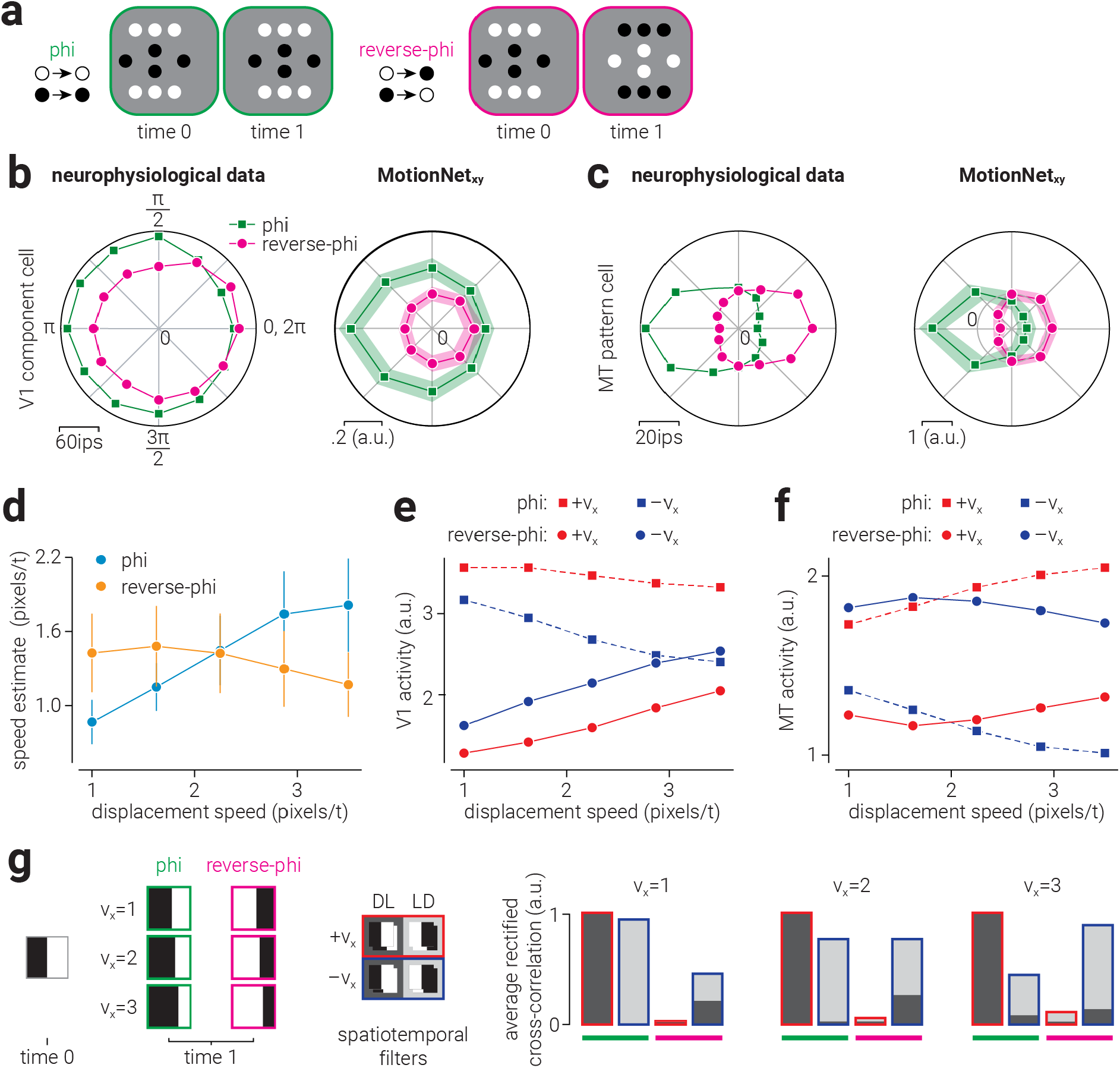
Biological and artificial visual system responses reverse-phi motion. (**a**) Illustration of phi and reverse-phi motion: between time zero and one all of the dots move to the right. For reverse-phi motion, the dots also reverse in polarity and are typically perceived as moving in the opposite direction. (**b-c**, left) Replotted data from Duijnhouwer and Krekelberg (2016) showing V1 component-motion and MT pattern-motion cell responses to standard- and phi-motion stimuli. (**b-c**, right) Average responses of MotionNet_xy_ V1 and MT units to equivalent stimuli. (**d**) Speed estimated by MotionNet_xy_ in response to phi and reverse-phi motion stimuli as a function of dot displacement speed. (**e-f**) The average activity of (**e**) V1 and (**f**) MT units that prefer motion in the direction of dot displacement (+vx) or the opposite direction (-vx), in response to phi and reverse-phi motion, as a function of displacement speed. Dashed and solid lines indicate response to phi and reverse-phi motion, respectively. (**g**) Illustration of simulation demonstrating misestimation of reverse-phi displacement. (**g**, left) Phi and reverse-phi edge stimuli with three different displacement distances are cross-correlated with four spatiotemporal filters tuned to rightward (+vx) and leftward (-vx) motion, with either dark-light (DL) or light-dark (LD) polarity arrangement. (**g**, right) Average rectified cross-correlation values, normalized to the maximum value. Stacked bars indicate the combined cross-correlation with opposite polarity filters tuned to the same direction. Rightward and leftward cross-correlation values are bordered in red and blue, respectively, and phi and reverse-phi results are underlined in green and magenta, respectively. Shaded areas in (**b-c**, right) and error bars in (**d**) indicate s.d. of average responses across 10 networks.

### Spatiotemporal tuning distributions and connections

In biological visual systems the tuning of spatiotemporal neurons in V1 and MT to spatial and temporal frequency follows a log-normal distribution (O’Keefe et al., 1998; Wang & Movshon, 2016; **Fig. 4a-d**, left). Similarly, we found that the preferred spatial and temporal frequencies of V1 and MT units in MotionNet_xy_ also followed a log-normal distribution (**Fig. 4a-d**, right). Speed is determined by the ratio of spatial and temporal frequency, meaning that different combinations of spatial and temporal frequencies could be used to achieve the same speed selectivity. For example, the same speed could be produced by a combination of low spatial and temporal frequency, or high spatial and temporal frequency. How might this be implemented in terms of the readout of V1 activity by speed-selective MT units? We established the preferred speed to which MotionNet_xy_’s MT units were tuned and separated these into “low” or “high” speed groups using a median split. We then compared the spatiotemporal tuning distributions of V1 units to which each group was maximally connected (**Fig. 4e-f**), i.e., weights with the highest positive values. We found that compared to MT units tuned to fast speeds, slow tuned units primarily received input from V1 units tuned to high spatial frequency and low temporal frequency. These results are consistent with work showing that the preferred speed of macaque MT neurons, as measured using dot motion stimuli, is negatively correlated with their preferred spatial frequency and positively correlated with their preferred temporal frequency (Priebe et al., 2003).

**Figure 4.**
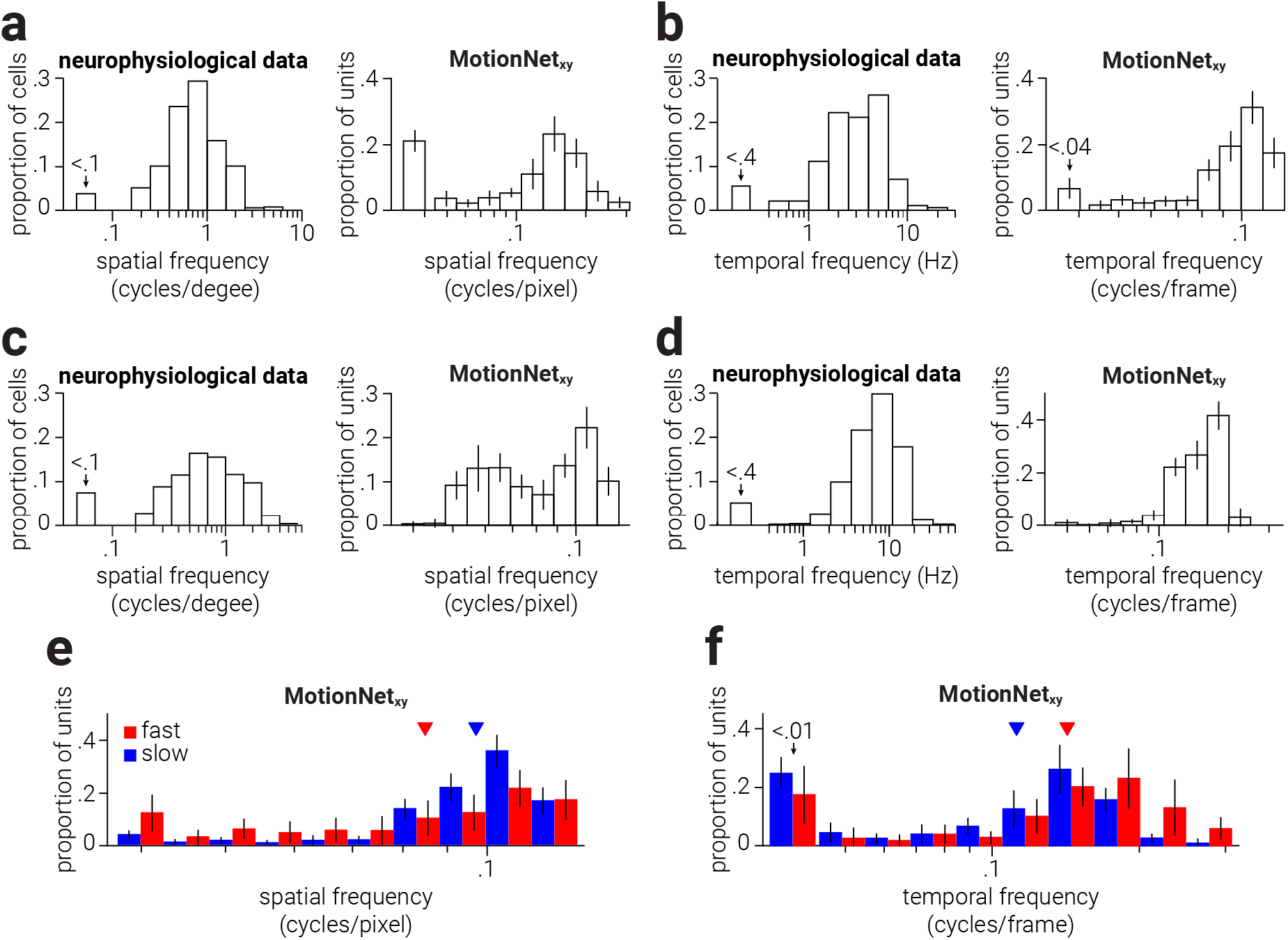
Biological and artificial visual system spatiotemporal tuning properties. (**a-b**, left) Replotted data from O’Keefe et al. (1998) showing the proportion of spatiotemporal cells in owl monkey V1 tuned to a range of spatial (**a**) and temporal frequencies (**b**). (**a-b**, right) Same as (a-b, left), but for V1 units of MotionNet_xy_. (**c-d**, left) Reanalysed data from Wang and Movshon (2016) showing the proportion of spatiotemporal cells in macaque MT tuned to a range of spatial (**c**) and temporal frequencies (**d**). (**c-d**, right) Same as (**c-d**, left), but for MT units of MotionNet_xy_. (**e-f**) Same as (**a-b**, right), but split into V1 units most strongly connected to MT units tuned to slower or faster speeds. Coloured arrows in (**e-f**) indicate the distributions means.

### Separable and covariate spatiotemporal tuning

Just as neurons can be classified according to their direction selectivity (i.e., component-/pattern-motion), they can classified by their spatiotemporal selectivity. In particular, neurophysiological evidence shows that V1 neurons are separately tuned to either spatial or temporal frequency. That is, they respond most strongly to a particular spatial frequency, regardless of the temporal frequency, or vice versa (Foster, Gaska, Nagler, & Pollen, 1985; Priebe, Lisberger, & Movshon, 2006; Tolhurst & Movshon, 1975). By contrast, some MT neurons are tuned to object speed, such that their sensitivity to spatial frequency is dependent on temporal frequency (Perrone & Thiele, 2001; Priebe et al., 2003). To identify whether a neuron has separable tuning or speed tuning, its response can be measured for a range of spatial and temporal frequencies. If the neuron has separable spatiotemporal tuning, the peak responses will align either horizontally or vertically with a particular spatial or temporal frequency (**Fig. 5a**, top). By comparison, if a neuron is tuned to speed, the peak responses will extend radially from the origin, with the angle indicating the speed to which the neuron is tuned (**Fig. 5a**, bottom). The fit of a two-dimensional Gaussian that is either aligned cardinally (horizontally/vertically) or radially to this activity can be used to quantitatively classify neurons as either separable or speed tuned (**Fig. 5a**, left). That is, in the same way as the response of a unit to plaid stimuli can be classified as component- or pattern-motion selective based on its alignment to the plaid vs. sinewave directions, we can use the radial vs. cardinal alignment of a unit’s responses to different spatial and temporal sinewaves to classify it as either separable- or speed-tuned. We performed this classification analysis on the V1 and MT units in MotionNet_xy_ and found that, in line with biological systems (Priebe et al., 2003), V1 units were separably tuned, while MT units showed a mixture of independent and speed tuning (**Fig. 5b**).

Just as is observed in macaque (**Fig. 5c**), we found a diverse range of MT units that were component-/pattern-motion selective and showed separable/speed tuning (**Fig. 5d**). It is possible that direction and speed selectivity properties are related among MT units, that is, a unit selective for complex direction (pattern-motion) may be more likely to be selective for complex speed. We tested this in MotionNet_xy_ found a positive correlation between pattern and speed indices of MT units (n=568, Pearson *r*=.72, *P*=1.9×10^-93^; **Fig. 5e**).

**Figure 5.**
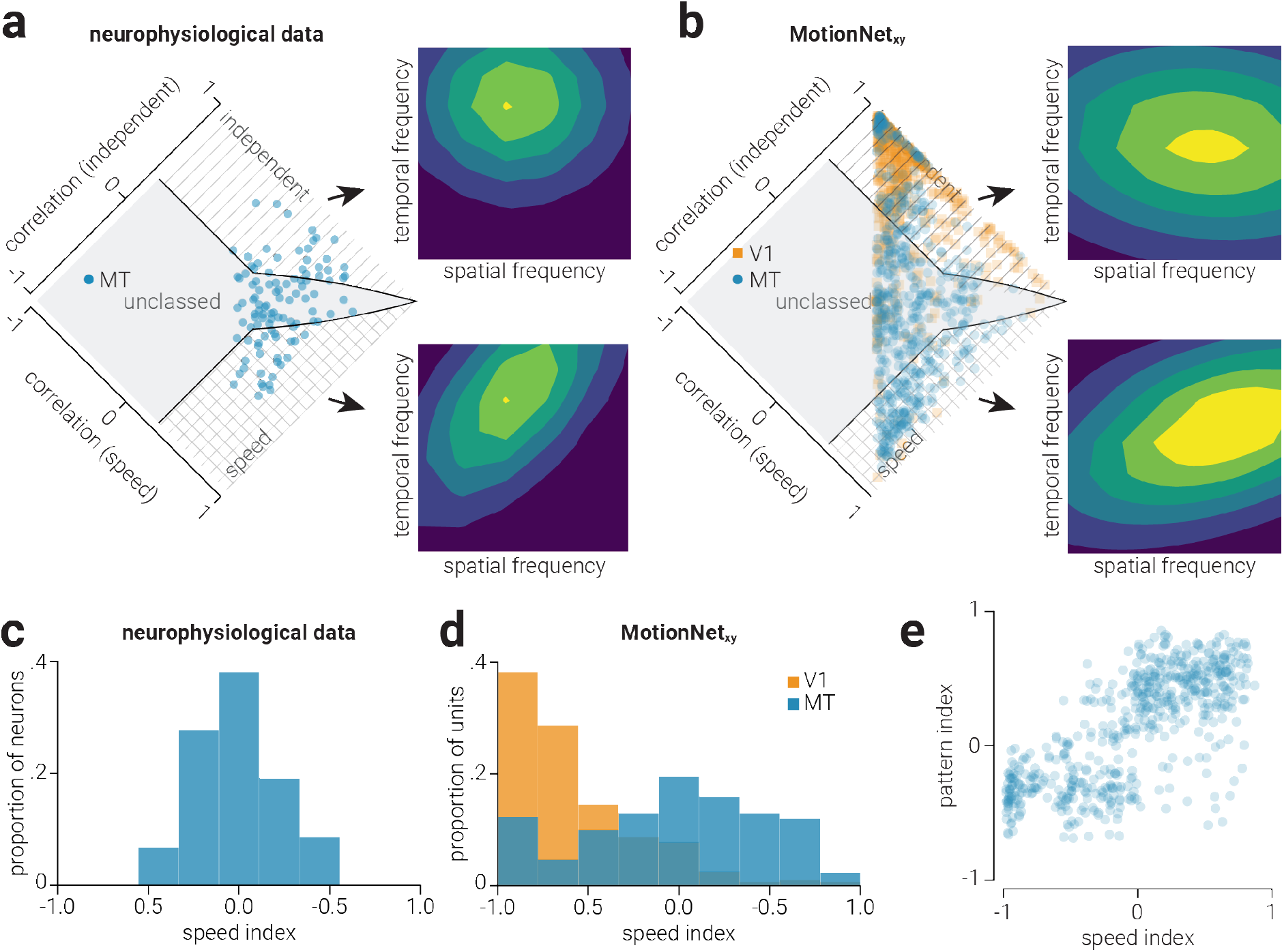
Biological and artificial visual system speed selectivity. (**a**) Data from (Priebe et al., 2003) showing responses of macaque MT neurons, that have separable (top) or speed (bottom) selectivity, to sinewave gratings at different spatial and temporal frequencies. The distribution plot shows the population of single neuron responses, and whether they are classified as separable or speed selective. (**b**) Same are (**a**) but for MotionNet_xy_ V1 and MT units; single polar plots (top and bottom) both show responses of MT units classified as either separate or speed selective. (**c-d**) Proportion of (**c**) macaque MT neurons (Priebe et al., 2003) and (**d**) MotionNet_xy_ V1 and MT units as a function of their speed index. (**e**) Scatter plot showing the relationship between speed and pattern selectivity for MotionNet_xy_ MT units.

### How do motion signals interfere with each other?

We next considered situations in which motion signals can degrade or may interfere with each other. First, we tested how the response to a moving dot pattern is affected by superimposing dots moving at different speeds. Biological visual systems exhibit inhibitory mechanisms that are thought to reduce noise and sharpen activity in response to visual features. For instance, experimenters have presented moving dot patterns and then overlaid dots moving in a different direction. V1 neurons are not substantially affected by this manipulation; however, MT neurons show *direction opponency* and are suppressed by dots moving in a non-preferred direction (Qian & Andersen, 1994; Rust, Mante, Simoncelli, & Movshon, 2006; Snowden, Treue, Erickson, & Andersen, 1991). We previously found comparable responses within a neural network trained to classify image velocity (Rideaux & Welchman, 2020). However, MT neurons also exhibit *speed opponency* and are suppressed by dots moving in a non-preferred speed (Mikami et al., 1986; **Fig. 6a-b**, left). We tested whether this noise reduction mechanism was also present in MotionNet_xy_ and found the same patterns of responses among MT units (**Fig. 6a-b**, right).

**Figure 6.**
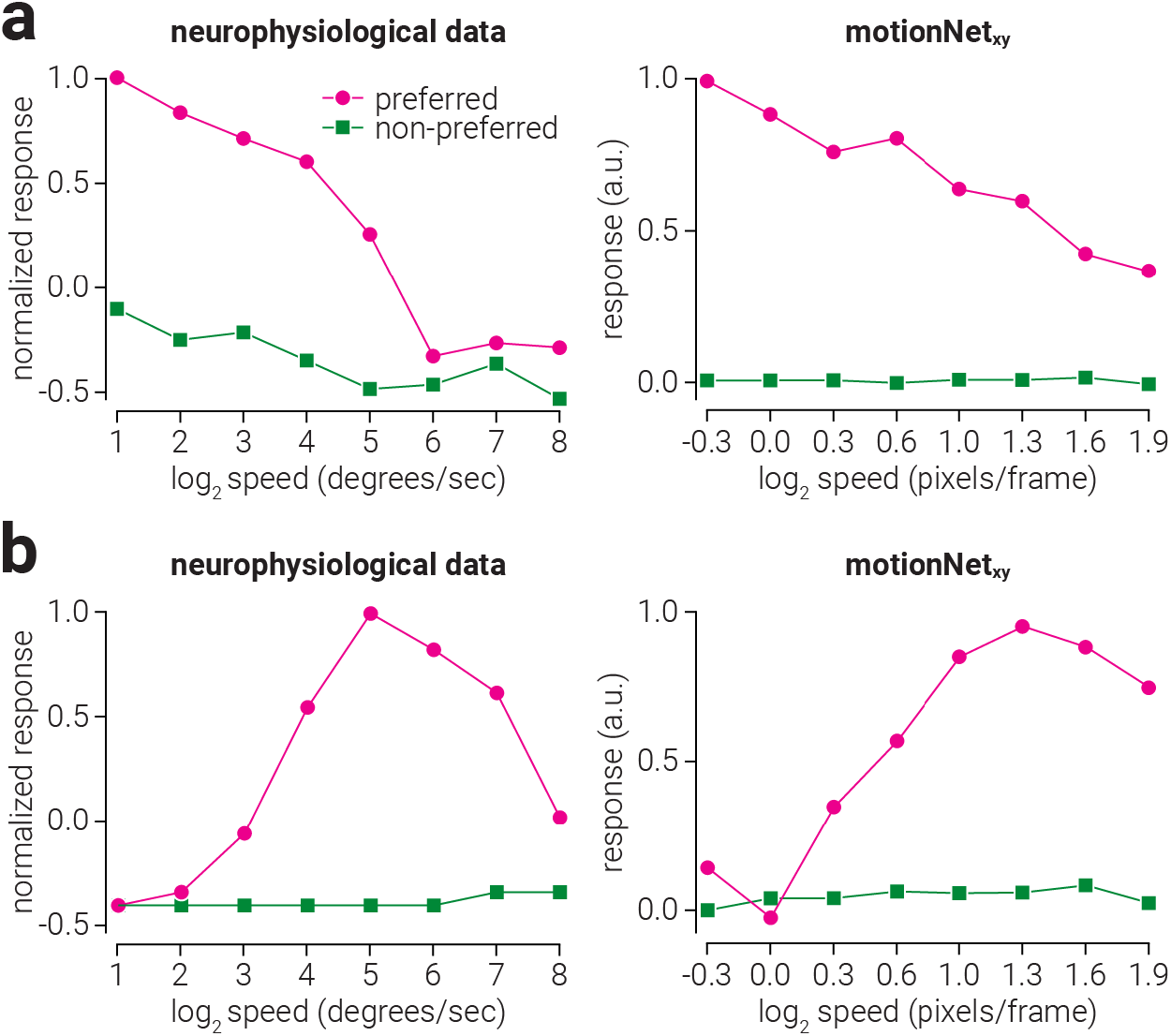
Biological and artificial visual system mechanisms of speed opponency. (**a-b**, left) Replotted data from Mikami, Newsome, and Wurtz (1986) showing the response of two MT neurons to a dot moving either in its preferred or non-preferred direction over a range of speeds. (**a-b**, right) The same as (**a-b**, left), but for the responses of selected MotionNet_xy_ MT units.

We then tested MotionNet_xy_ with random dot stimuli that have been widely used to study motion. Using these stimuli it is possible to precisely titrate the relationship between dots moving in a particular direction (the signal) and dots moving in a randomly chosen direction (noise). We tested the ability of MotionNet_xy_ to correctly estimate the direction of motion by varying the proportion of signal and noise dots in the stimulus (**Fig. 7a**). Like individual neuronal responses (Britten, Shadlen, Newsome, & Movshon, 1992; **Fig. 7b**) and macaque monkey psychophysical judgments (**Fig. 7c**, blue markers), we found graceful degradation in estimates of motion direction (**Fig. 7c**, red markers). We showed that reducing motion coherence reduces the accuracy of direction estimates, but how are speed judgements influenced? Previous psychophysical evidence shows that humans underestimate the speed of dot motion with reduced coherence (Schütz et al., 2010; **Fig. 7d**, orange markers). We tested how MotionNet_xy_ estimated the speed of dot motion at different coherence levels and found the same pattern of results (**Fig. 7d**, cyan markers).

As the directions of noise dots are uniformly distributed around 360°, the average velocity of the noise is zero. The underestimation of the speed of partially coherent dot motion stimuli appears to adhere to a linear trend that is equal to the weighted average of noise (zero) and signal (non-zero) speed, where the weights are equal to the proportion of noise and signal dots. Thus, a possible explanation for this bias is that it is produced by pooling of noise and signal by the network. We reasoned that if the bias is produced by pooling of noise and signal, then we would expect that the response of the network to 50% coherence motion to be similar to the pooled responses to 0% and 100% coherence.

Consistent with this explanation, we found that the average activity of MT units in response to 50% coherence motion could be predicted with high accuracy by averaging their responses to 0% and 100% coherence (n=64, Pearson *r*=.99, *P*=1.4×10^-50^; **Fig. 7e**).

**Figure 7.**
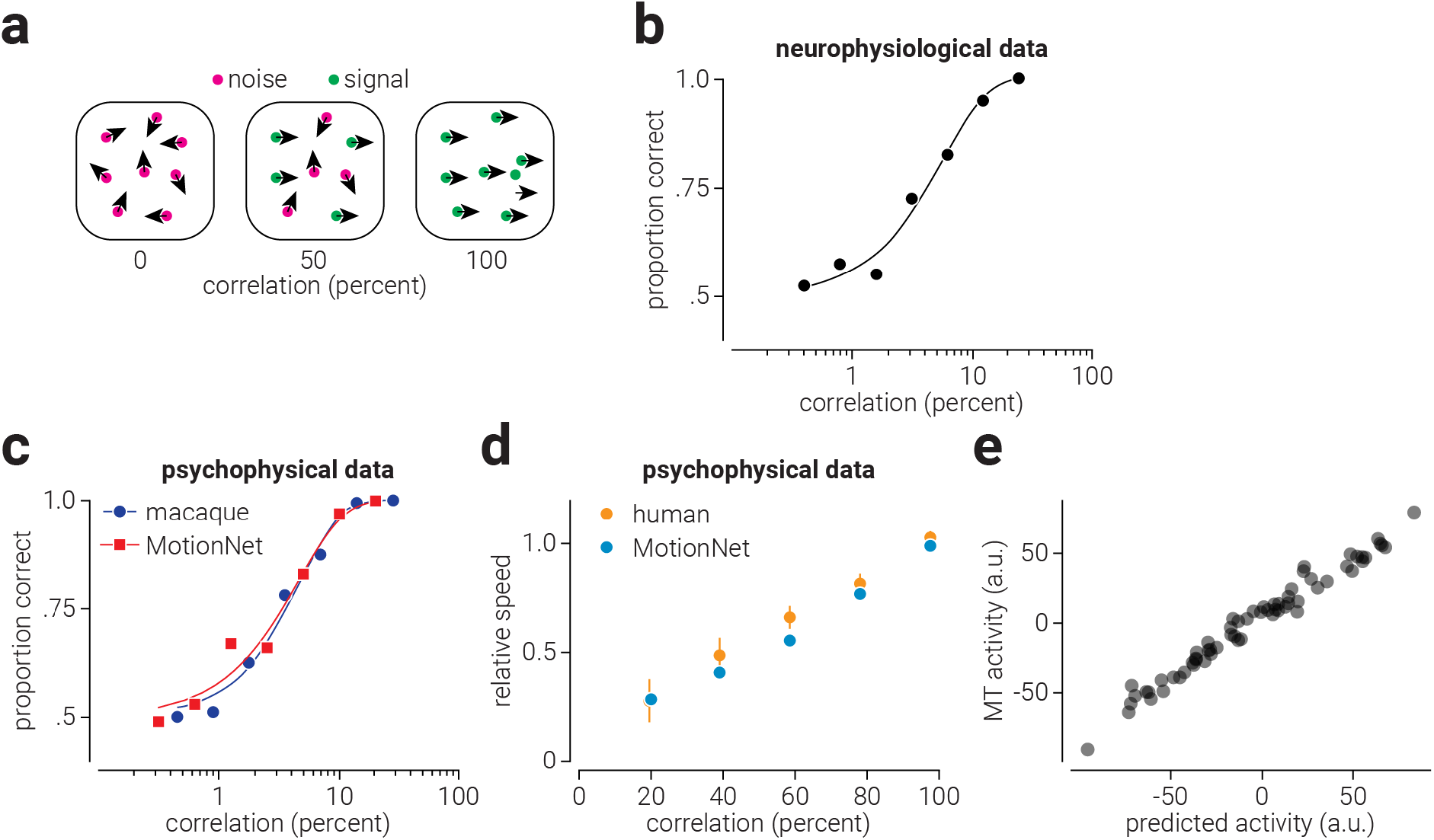
Biological and artificial visual system responses to reduced motion coherence. (a) Illustration of global dot motion stimulus at varying degrees of motion signal correlation. (b) Replotted data from Britten et al. (1992) showing a neurometric function that describes the sensitivity of an MT neuron to motion signals of increasing correlation. (c) Same as (b) but for psychophysical performance of a macaque (and MotionNet_xy_) on a direction discrimination task. The blue circles and red squares indicate mean performance and the corresponding coloured lines show the Weibull function used to determine threshold performance. (d) Replotted data from (Schütz et al., 2010) showing human speed estimates, relative to the signal speed, of dot motion stimuli at different levels of signal correlation. The blue dots in (d) show MotionNet_xy_’s relative speed estimates for similar stimuli. (e) The activity of MotionNet_xy_’s MT units in response to a 50% coherence motion stimulus, as a function of the activity predicted by averaging their responses to 0% and 100% coherence stimuli.

## DISCUSSION

The ability to see movement underpins adaptive behaviours ranging from depth estimation to navigation and grasping. Here we explore and explain the neural computations that support motion estimation in biological systems by investigating the structures that emerge in an artificial system trained to estimate the velocity of image sequences. Using complete access to the artificial system, we reveal aspects of the neural architecture that instantiates the motion estimation, producing concrete predictions for future empirical study. Specifically, we show that (i) the network overestimates the speed of slow reverse-phi motion while underestimating the speed of fast reverse-phi motion because of the correlation between reverse-phi motion and the spatiotemporal receptive fields tuned to motion in opposite directions, (ii) compared to MT units tuned to fast speeds, those tuned to slow speeds primarily receive input from V1 units tuned to high spatial frequency and low temporal frequency, (iii) there is a positive correlation between the pattern-motion and speed selectivity of MT units, and (iv) the network recapitulates human underestimation of low coherence motion stimuli, which is explained by pooing of noise and signal motion.

Reverse-phi motion is perceived as moving in the opposite direction to the actual movement (Anstis, 1970). The manner in which this image manipulation influences the preferred direction of neurons and the perceived direction of movement has been documented (Duijnhouwer & Krekelberg, 2016). Here we show that in addition to these effects related to direction, this manipulation may also produce biases in perceived speed. Further, we lay bare the computational mechanism explaining this new phenomenon. That is, the similarity between reverse-phi motion and receptive fields of spatiotemporal units tuned to opposite velocities. While some behavioural evidence for this bias has previously been documented (Parthasarathy, 2019; Ruda et al., 2016), future psychophysical and neurophysiological work is needed to directly test these predictions.

We previously showed that multiple physiological and psychophysical phenomena in motion processing are recapitulated by a network trained to classify the velocity of moving image sequences (Rideaux & Welchman, 2020). For example, we found that the anisotropic distribution of direction preferences in units in a layer representing V1 matched that of neurons in mouse V1. Here we found that the distribution of spatial and temporal frequency tuning also matched that found in macaque V1 and MT (i.e., log-normal distribution of neuronal frequency preference). Previous electrophysiological work suggested that the MT neurons tuned to low speeds primarily receive input from V1 neurons tuned to high spatial frequency and low temporal frequency, whereas the opposite pattern of transmission was true for MT neurons tuned to high speed (Priebe et al., 2003). This evidence was based on the activity of MT neurons, as measuring connections and preferences of neurons across cortical regions on a sufficiently large scale is beyond the limitations of current biological techniques. By contrast, this analysis is made possible within the artificial system, and we find evidence consistent with previous hypotheses: slow-tuned MT units receive more input from high spatial and low temporal frequency V1 units than fast-tuned MT units.

Considerable work has been undertaken to understand how the properties of spatiotemporal neurons in MT are distinguished from those in V1, as this knowledge can provide insight into the hierarchical computations that underlie motion processing. Neurons can be classified by their direction selectivity (i.e., component-/pattern-motion) or spatiotemporal selectivity (i.e., separate/speed). V1 only contains neurons selective for component-motion and separate spatiotemporal frequencies, while neurons selective for pattern-motion and speed are found in MT. This dichotomy supports the notion that “simple” motion signals from V1 are pooled in MT, yielding selectivity for more “complex” signals. However, neurophysiological work shows that the selectivity of many MT neurons is indistinguishable from those in V1. We found the same pattern of results for MotionNet_xy_: the MT layer comprised a mixture of units tuned to component- and pattern-motion, and separate spatiotemporal frequency and speed.

Our results indicate that rather than MT units either being separately tuned to a particular spatial/temporal frequency or speed, the distribution of speed selectivity in MT reflected a continuum along this dimension. This tuning diversity is consistent with neurophysiological evidence from macaque (Priebe et al., 2003). We also found a positive relationship between direction and speed selectivity of MT units, indicating that units tuned to complex motion signals in one domain (e.g., direction) were more likely to be tuned to complex signals in the other (e.g., spatiotemporal). Given that the complexity of the selectivity for both direction and speed is derived from the same characteristic, i.e., diversity of connection weights between V1 and MT, it seems reasonable to expect that these properties would be related. However, in contrast, previous neurophysiological work did not find evidence for this relationship in macaque (Priebe et al., 2003). A possible explanation for this conflict is that there was an insufficient range of speed selectivity in the neurophysiological sample to detect the relationship. In our data, we recorded units ranging almost the entire speed selectivity continuum, whereas the neurophysiological data accounted for approximately half this range (possibly due to noise within the biological system reducing the effectiveness of the classification technique). More neurophysiological work is needed to test this possibility.

We previously demonstrated that the tendency for humans to underestimate the speed of objects moving at low visibility could be explained by the lawful relationship between spatiotemporal contrast and speed in natural image sequences, rather than exposure to a non-uniform distribution of motion speeds in the environment, i.e., the “slow-world” bias (Rideaux & Welchman, 2020). There have been multiple psychophysical demonstrations of the bias under conditions of reduced contrast (Hürlimann, Kiper, & Carandini, 2002; Sotiropoulos, Seitz, & Seriès, 2014; Vintch & Gardner, 2014; Weiss, Simoncelli, & Adelson, 2002); however, there is also evidence that humans underestimate the speed of dot motion stimuli with reduced signal coherence (Schütz et al., 2010). This could be interpreted as evidence for the slow-world account, as reducing signal coherence likely reduces estimation certainty. However, we tested MotionNet_xy_ and found the same pattern of results: the network underestimated the speed of dot motion stimuli with reduced signal coherence. Importantly, this phenomenon was an outcome of pooling signal and noise together, and unrelated to the mechanism that produces underestimation of low contrast motion signals.

In recent years, deep neural networks comprising many layers have surpassed the human performance on many tasks, e.g., object recognition (He, Zhang, Ren, & Sun, 2016; Russakovsky et al., 2015). However, their scale and complexity often obscures inspection; limiting understanding of their internal processes as much as in biological systems. Here, we constrain the size of the artificial system, allowing us to apply *in silico* electrophysiological techniques that lay bare and understand the processes that underlie perceptual (mis-)estimation of velocity. We demonstrate how optimising motion estimation in an artificial network using natural images recapitulates a diverse array of neurophysiological and perceptual phenomena. More importantly, we use this technique to explain the computational basis of existing perceptual phenomena and generate predictions for some yet to be tested.

## Acknowledgements

We thank Dr. Parthasarathy for their insights relating to the perceived speed of reverse-phi stimuli. The work was supported by the Leverhulme Trust (ECF-2017-573) and the Issac Newton Trust (17.08(o)).

